# PANOMIQ: A Unified Approach to Whole-Genome, Exome, and Microbiome Data Analysis

**DOI:** 10.1101/2024.09.17.613203

**Authors:** Shivani Srivastava, Saba Ehsaan, Linkon Chowdhury, Muhammad Omar Faruk, Abhishek Singh, Anmol Kapoor, Sidharth Bhinder, M. P. Singh, Divya Mishra

## Abstract

The integration of whole-genome sequencing (WGS), whole-exome sequencing (WES), and microbiome analysis has become essential for advancing our understanding of complex biological systems. However, the fragmented nature of current analytical tools often complicates the process, leading to inefficiencies and potential data loss. To address this challenge, we present PANOMIQ, a comprehensive software solution that unifies the analysis of WGS, WES, and microbiome data into a single, streamlined pipeline. PANOMIQ is designed to facilitate the entire analysis process from raw data to interpretable results. It is the fastest algorithm that can achieve results much more quickly compared to traditional pipeline approaches of WGS and WES analysis. It incorporates advanced algorithms for high-accuracy variant calling in both WGS and WES, along with robust tools for characterizing microbial communities. The software’s modular architecture allows for seamless integration of these diverse data types, enabling researchers to uncover complex interactions between host genomics and microbiomes. In this study, we demonstrate the capabilities of PANOMIQ by applying it to a series of datasets encompassing a wide range of applications, including disease association studies and environmental microbiome profiling. Our results highlight PANOMIQ’s ability to deliver comprehensive insights, significantly reducing the time and computational resources required for multi-omic analysis. By providing a unified platform for WGS, WES, and microbiome analysis, PANOMIQ offers a powerful tool for researchers aiming to explore the full spectrum of genomic and microbial diversity. This software not only simplifies the analytical workflow but also enhances the depth of biological interpretation, paving the way for more integrated and holistic studies in genomics and microbiology.

## Introduction

Advances in genomic technologies have revolutionized our understanding of human biology, health, and disease (Park, S. T., & Kim, J. 2016). Among these, whole-genome sequencing (WGS) provides comprehensive insights into the entire genetic makeup of an organism, while exome sequencing focuses on the protein-coding regions, which are often the most relevant to disease. Simultaneously, the study of the human microbiome has uncovered the crucial role that microbial communities play in maintaining health and contributing to disease states (Ormond et al., 2010). Each of these approaches—WGS, exome sequencing, and microbiome analysis—offers unique and valuable perspectives, yet they also generate vast and complex datasets that pose significant analytical challenges (Brlek et al., 2024).

Integrating these diverse data types is essential for a more holistic understanding of biological processes and disease mechanisms. However, the heterogeneity, scale, and complexity of genomic and microbiome data make such integration difficult. Traditional tools often focus on analyzing one type of data in isolation, resulting in fragmented insights that fail to capture the full picture (Bollas et al., 2024). The lack of a unified approach to analyze whole-genome, exome, and microbiome data simultaneously is a major bottleneck in the field, limiting the potential for comprehensive discoveries (Olson et al., 2023).

To address this critical gap, we present PANOMIQ—a novel, unified tool designed to seamlessly integrate and analyze whole-genome, exome (Fig.1), and microbiome data. PANOMIQ leverages cutting-edge algorithms to manage the scale and complexity of these datasets, providing a coherent analytical framework that can extract meaningful insights across all three domains. This unified approach not only enhances the accuracy and efficiency of genomic analyses but also opens new avenues for understanding the complex interactions between the human genome and its associated microbiome.

**Fig. 1:**
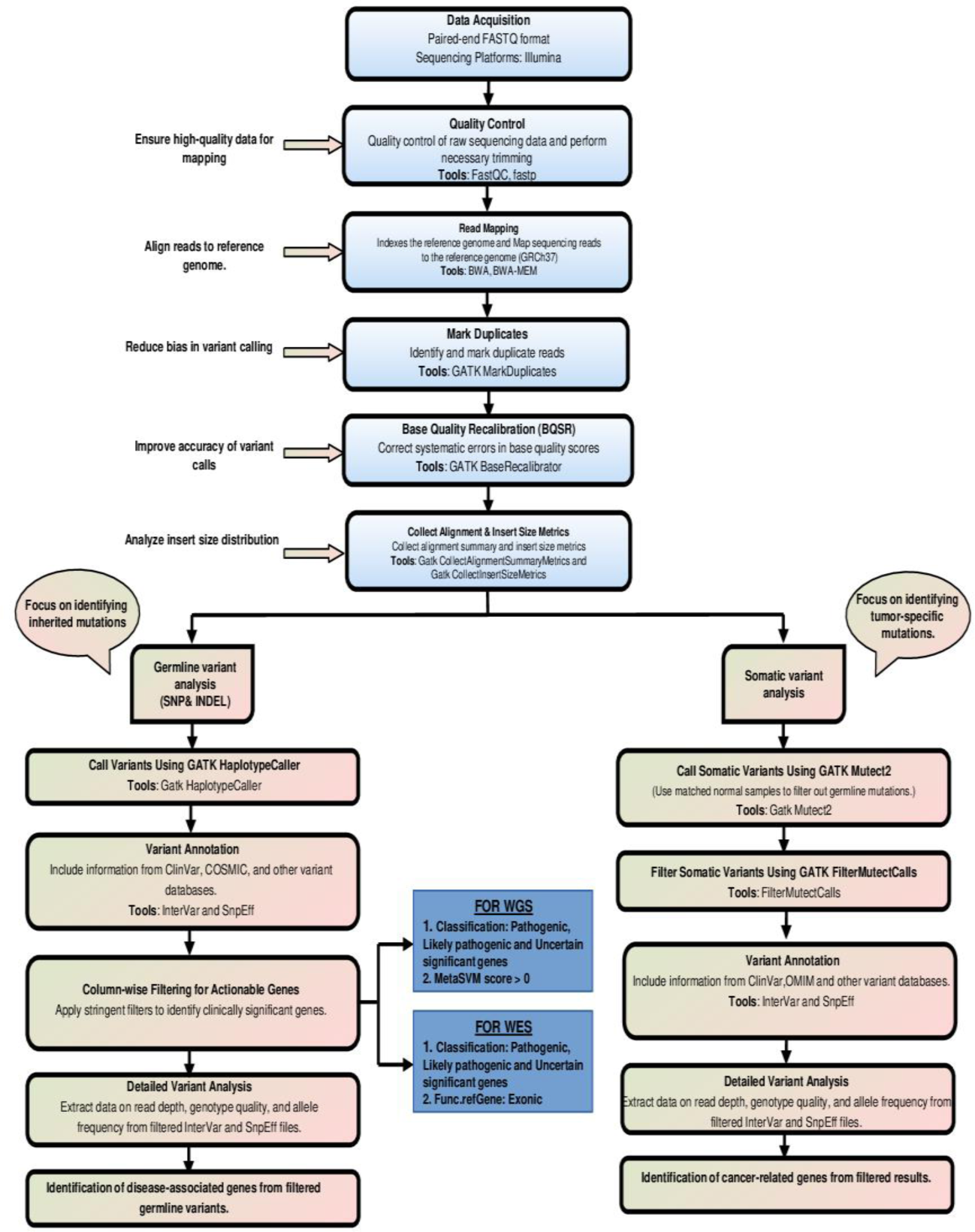
Workflow for Whole Genome Sequencing (WGS) and Whole Exome Sequencing (WES) Analysis Automated in PANOMIQ Solution: This figure provides a comprehensive overview of the automated analysis workflow for Whole Genome Sequencing (WGS) and Whole Exome Sequencing (WES) within the PANOMIQ solution. The workflow is designed to streamline and optimize genomic analysis, ensuring efficiency and accuracy throughout the process.

The implications of PANOMIQ are far-reaching. For researchers, it offers a powerful tool to explore the intricate relationships between genetic variations and microbiome composition, which could lead to breakthroughs in the study of complex diseases. Clinicians could leverage this integrated analysis to improve personalized medicine, tailoring treatments based on a comprehensive understanding of a patient’s genomic and microbiomic profile. Furthermore, PANOMIQ contributes to the broader field of bioinformatics by setting a new standard for multi-omic data integration and analysis.

In this study, we aim to develop, validate, and demonstrate the effectiveness of PANOMIQ across various use cases, showcasing its potential to transform both research and clinical practice. The following sections of this paper will detail the methodology behind PANOMIQ, present validation results, and discuss its application in real-world scenarios.

## Materials and Methods

### 1. Whole Exome Sequencing (WES) Analysis

#### 1.1. Primary Analysis

##### Preprocessing and Quality Assessment

Paired-end FASTQ files were downloaded from the NCBI-Sequence Read Archive (SRA) and subjected to quality assessment using FastQC (Andrews, 2010). Critical parameters such as per base sequence quality, per sequence GC content, and adapter content were evaluated to ensure data integrity. Low-quality reads and adapter sequences were filtered out using Fastp (version 0.20.0) (Chen et al., 2018). Reads with more than 10% N bases, more than 50% bases with quality scores below 5, or an average quality score below 10 were removed (Yang et al., 2023).

##### Reference Genome Preparation

The GRCh37 reference genome was downloaded from the Ensembl database (Cunningham et al., 2015). The genome was decompressed and indexed using Samtools (Li et al., 2009; Li, 2011), and a sequence dictionary was created with GATK (McKenna et al., 2010) to facilitate efficient access during alignment and variant calling.

##### Mapping and BAM File Processing

Sequencing reads were aligned to the GRCh37 reference genome using the BWA-MEM algorithm (Li & Durbin, 2009), producing BAM files for each sample. The BAM files were then processed using GATK tools to mark duplicate reads and perform Base Quality Score Recalibration (BQSR) (Van et al., 2013). This step corrected for systematic biases introduced during sequencing, enhancing the accuracy of downstream variant calling.

#### 1.2. Secondary Analysis

##### Variant Calling

Germline variants were identified using GATK HaplotypeCaller (Van der Auwera & O’Connor, 2020). The tool constructs haplotypes from aligned reads and calls variants relative to the reference genome, producing a VCF file with SNPs and indels. For somatic variant detection, GATK Mutect2 (Benjamin et al., 2019) was used, particularly focusing on low-frequency variants in exonic regions relevant to cancer research.

#### 1.3. Tertiary Analysis

##### Variant Annotation

Variant annotation was performed to predict the potential impact of genetic variants on gene function. InterVar (Li & Wang, 2017) was used to classify variants based on the American College of Medical Genetics and Genomics (ACMG) guidelines, providing clinical significance ratings. Additionally, SnpEff (Cingolani et al., 2012) was employed to predict the effects of SNPs and indels on protein coding sequences, including the identification of missense, nonsense, and splicing mutations. This combined approach facilitated a comprehensive understanding of the clinical relevance of the variants detected, essential for interpreting genetic disorders (Fig. 2).

**Fig. 2:**
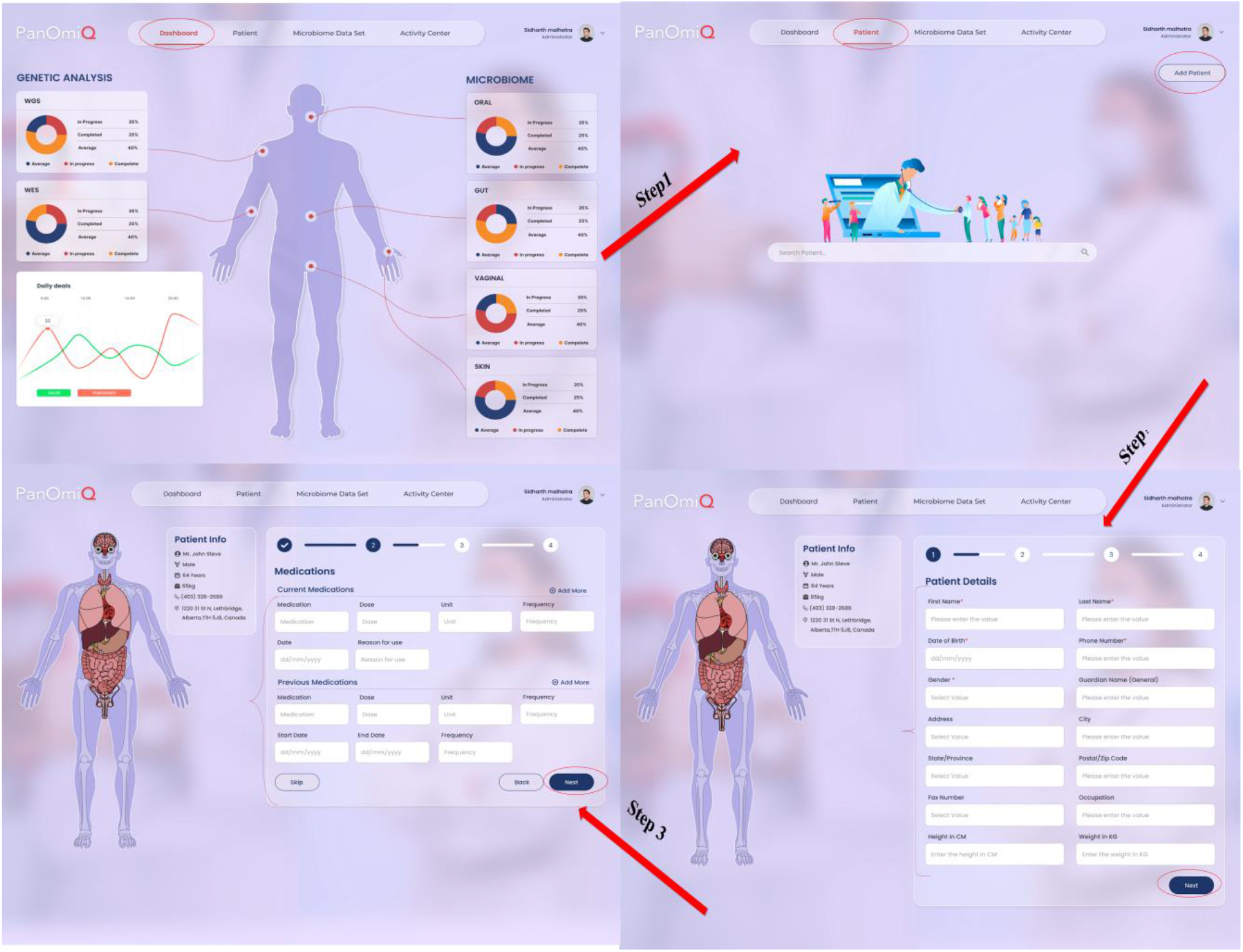
PANOMIQ Front-End Interface for Primary Analysis: This figure illustrates the PANOMIQ front-end interface used for initiating primary analysis. It demonstrates how users can upload FASTQ files and enter individual details for genomic analysis. Key elements include: **FASTQ File Upload**: The section where users upload raw sequencing data files in FASTQ format. **Individual Details Entry**: The interface for entering and managing metadata associated with each sample, including patient identification and demographic information. This interface facilitates the seamless integration of raw data with sample-specific details, setting the stage for the subsequent processing and analysis phases.

### 2. Whole Genome Sequencing (WGS) Analysis

#### 2.1. Primary Analysis

##### Preprocessing and Quality Assessment

FASTQ files obtained from whole-genome sequencing were initially assessed using FastQC for quality control. Subsequent filtering of low-quality reads and adapter trimming was performed using Fastp (Chen et al., 2018). The filtering criteria included removal of reads with more than 10% N bases and bases with quality scores below 10.

##### Reference Genome and Known Sites Preparation

The GRCh37 reference genome was prepared and indexed as described for WES. Additionally, VCF files containing known sites of genetic variation were downloaded from Ensembl for use in BQSR (Van et al., 2013).

##### Mapping and BAM File Processing

Paired-end reads were aligned to the GRCh37 reference genome using BWA-MEM, followed by processing with GATK tools to mark duplicates and perform BQSR (Van et al., 2013). These steps ensured high-quality, reliable data for variant calling.

#### 2.2. Secondary Analysis

##### Variant Calling

Germline variant detection was carried out using GATK HaplotypeCaller (Van der Auwera & O’Connor, 2020), while somatic variants were identified using GATK Mutect2 (Benjamin et al., 2019). This approach allowed for comprehensive detection of both common and rare variants across the entire genome.

#### 2.3. Tertiary Analysis

##### Variant Annotation

Variant annotation for WGS data was conducted using InterVar (Li & Wang, 2017) and SnpEff (Cingolani et al., 2012). InterVar provided clinical classifications according to ACMG guidelines, while SnpEff offered detailed predictions on the functional impact of genetic variants. This annotation process is crucial for interpreting the potential effects of genetic variations across the entire genome, particularly in understanding complex genetic disorders and guiding clinical decision-making.

### 3. Microbiome Analysis

#### 3.1. Primary Analysis

##### Initial Preprocessing

Raw sequencing data were preprocessed using Cutadapt (Martin, 2011) to remove adapter sequences and low-quality bases. This step ensured that only high-quality reads proceeded to downstream analysis. Further quality control and decontamination were performed using Kneaddata (Beghini et al., 2021), which removes host DNA contamination and filters low-quality reads, ensuring that the final dataset predominantly contains microbial sequences.

#### 3.2. Secondary Analysis

##### Taxonomic Classification

The filtered reads were subjected to taxonomic classification using Kraken (Wood & Salzberg, 2014; Breitwieser et al., 2019). Kraken’s k-mer based approach was employed to assign taxonomic labels, with a confidence threshold of 0.1 to reduce false positives. To improve the accuracy of taxonomic abundance estimates, Bracken (Lu et al., 2017) was used to refine the initial classifications, providing a more accurate representation of the microbial community composition (Fig. 3).

**Fig. 3:**
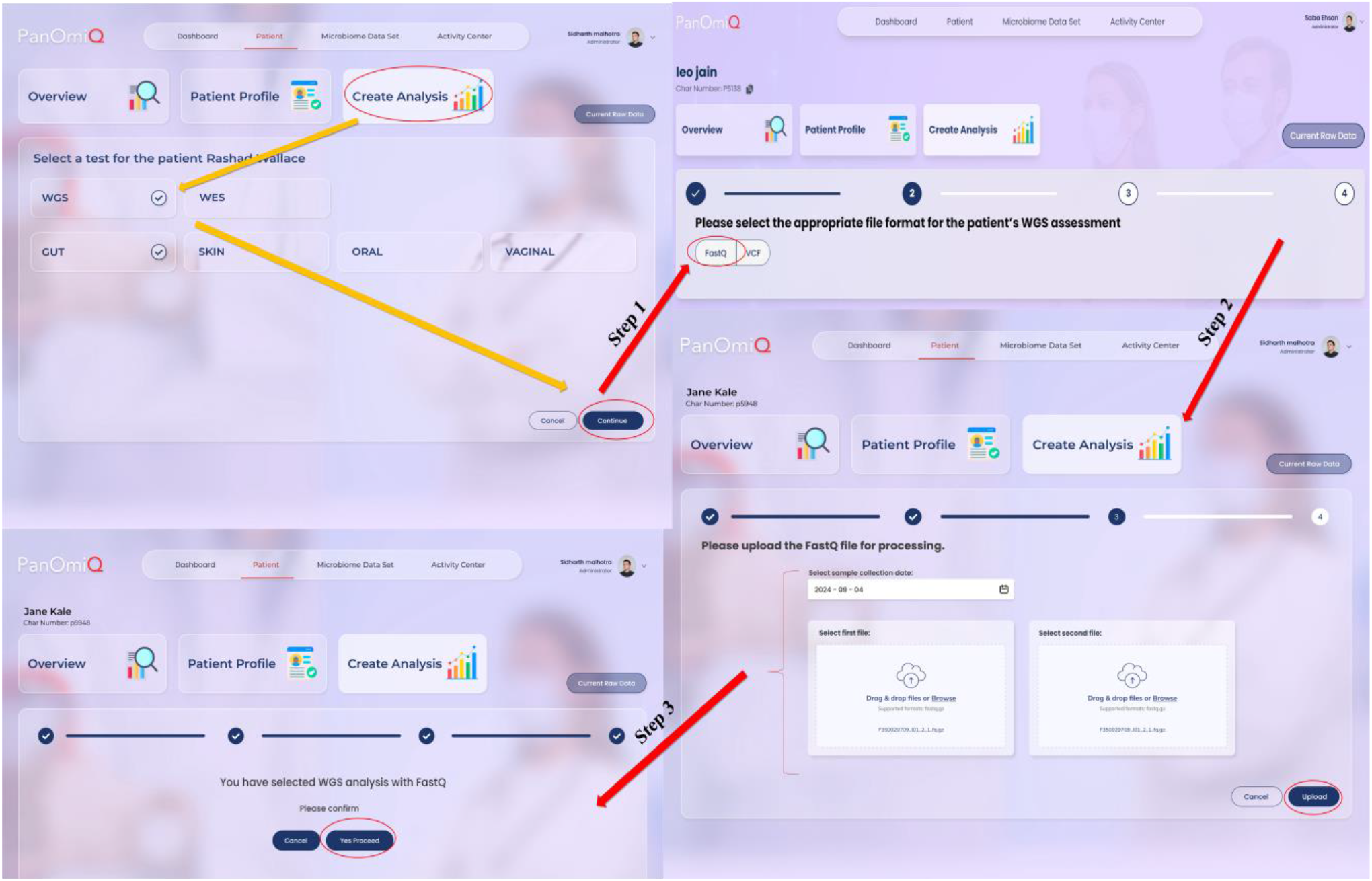
PANOMIQ Interface for File Upload and Option Selection: This figure illustrates the PANOMIQ interface, showcasing the available options for configuring analysis settings and uploading files to the server. Key features highlighted include: **Option Selection Panel:** Displays the various analysis options and parameters that users can choose to customize their workflow. **File Upload Section:** Demonstrates the process of uploading files to the server, including supported file formats and the steps involved in ensuring successful data transfer.The interface is designed to streamline the setup of analysis workflows and facilitate efficient data management within the PANOMIQ system.

#### 3.3. Tertiary Analysis

##### Visualization with Krona

For taxonomic data visualization, Krona was employed to generate interactive pie charts (Ondov et al., 2011). These charts provide a hierarchical view of the microbial community, allowing for detailed exploration of different taxonomic levels. Krona’s interactive features enable an intuitive examination of relative abundances from phylum to species level (Ondov et al., 2011).

##### Gut Microbiome Diversity Analysis

The processed taxonomic data were further analyzed for microbial diversity using Kraken Tools alpha_diversity (Lu et al., 2022). The Shannon diversity index, which measures species richness and evenness, was calculated (Shannon, 1948). For instance, a Shannon diversity index (SDIV) value of 3.174 was obtained, indicating a healthy microbial diversity within the gut microbiome. Additionally, the Firmicutes/Bacteroidetes (F/B) ratio, a key biomarker for gut health, was calculated, yielding a value of 3.1219. This higher F/B ratio suggests a greater abundance of Firmicutes, which may be indicative of certain health conditions or dysbiosis (Cummings et al., 1987; Turnbaugh et al., 2009; Friedman & Alm, 2012).

##### Final Table Generation in Genetic Variant Analysis

In genetic variant analysis, the final table generation is a critical step that consolidates and filters variant data for reporting or further analysis. This process integrates the outputs of variant calling algorithms, annotates variants with details such as genomic location and predicted impact, and applies stringent filters to retain only high-quality variants. Clinical interpretation contextualizes these variants against databases and literature, while quality control procedures eliminate artifacts and false positives. The resultant table supports clinical decision-making, research, and the dissemination of genomic findings (Fig. 4).

**Fig. 4:**
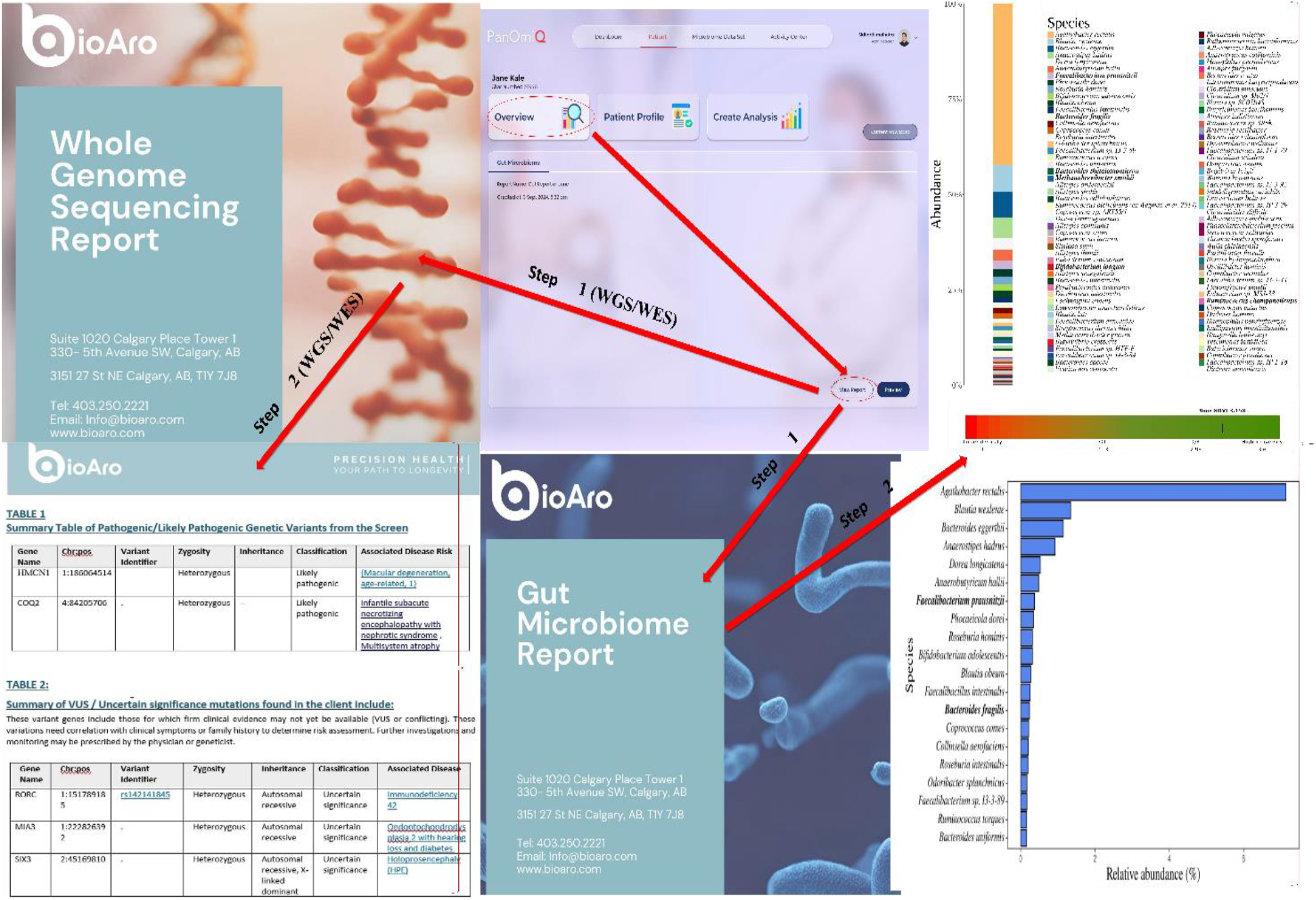
Cover Page of PANOMIQ Reports: This figure illustrates the cover page of the PANOMIQ reports, detailing its layout and design. The cover page includes: Report Title and Identification: Displays the title of the report and relevant identification details for easy reference. Summary Information: Provides a snapshot of key findings and analysis highlights. Download Instructions: Shows the available options for users to download the report, ensuring accessibility and convenience. The cover page is designed to provide a clear and professional overview of the report’s content, with easy access for downloading and review.

##### Data Filtering in WGS and WES

###### WGS Data Filtering

For Whole Genome Sequencing (WGS), filtering begins with variants of uncertain significance, as indicated by the InterVar column. Filters are applied based on MetaSVM scores (>0), which predict deleterious effects, and on pathogenicity, focusing on variants classified as “Pathogenic” or “Likely pathogenic.” Selected columns include basic genomic coordinates, gene annotations, and clinical significance data (e.g., ClinVar, OMIM, Orphanet). This process optimizes the extraction of clinically relevant data for downstream analysis.

###### WES Data Filtering

In Whole Exome Sequencing (WES), similar filtering processes are applied, focusing on variants within exonic regions (using the Func.refGene column) and those annotated in InterVar. The final table includes selected columns similar to WGS, ensuring the relevance and clinical utility of the filtered data.

###### Column Selection and Data Integration

Column-wise filtering in both WGS and WES targets key attributes like quality scores, allele frequencies, and functional impacts to identify clinically actionable genes. Essential columns include genomic coordinates, gene annotations, pathogenicity predictions, and links to databases like ClinVar, OMIM, and Orphanet. Read depth, genotype qualities, and allele frequencies are also extracted to assess variant reliability.

###### Finalizing the Table

The final table categorizes variants based on clinical significance, distinguishing between those requiring further validation and those with strong evidence of pathogenicity. The table is formatted for clarity and practical use, making it a valuable resource for clinicians, genetic counselors, and researchers in precision medicine and genetic research.

## Results: Microbiome Analysis

### 1. Stacked Bar Graph of Microbial Species Distribution

The stacked bar graph effectively illustrates the distribution and diversity of microbial species within the gut microbiome sample. Each color block in the graph corresponds to a specific microbial species, with keystone species highlighted in bold in the accompanying species list. These keystone species play a critical role in maintaining the stability and health of the gut ecosystem. Their presence and relative abundance can have significant implications for overall gut health. The species list includes key microbial species identified in the sample, providing a comprehensive overview of the gut microbiome composition (Segata et al., 2012; Gevers et al., 2012; Pasolli et al., 2019).

### 2. Gut Microbiome Diversity Visualization

Gut microbiome diversity, a key indicator of a balanced and healthy gut ecosystem, was visualized using the Shannon Diversity Index (SDIV). The diversity of microbial species is vital for supporting a range of functions, including digestion, immune response, and protection against pathogens (Lozupone et al., 2012). The SDIV for the gut microbiome in this study was calculated to be 3.174, which falls within the healthy reference range of 2.13 to 3.6 (Arumugam et al., 2011). This high level of diversity is indicative of a robust gut microbiome, which is generally associated with numerous health benefits such as enhanced immune function, better digestion, and a reduced risk of diseases like diabetes, inflammatory bowel disease (IBD), obesity, and liver diseases (Foster & Neufeld, 2013; Belkaid & Hand, 2014). Maintaining such diversity is crucial for overall well-being (David et al., 2014).

### 3. Visualization of Abundant Members of the Gut Microbiome

The relative abundance of the top 20 microbial species in the gut microbiome was visualized using a bar chart. This chart highlights the most and least abundant species, providing insights into the overall composition of the gut microbiome (Turnbaugh et al., 2006; Lynch & Pedersen, 2016). The length of each bar corresponds to the relative abundance of each species, allowing for an easy comparison across species. This type of visualization is essential for identifying key species and understanding their role in gut health, which is crucial for microbiome studies and related health research (Sender et al., 2016).

### 4. Keystone Bacteria in Gut Health

Certain keystone bacteria were identified with higher relative abundance in the gut microbiome sample compared to a healthy reference cohort. Keystone species like *Ruminococcus bicirculans, Eubacterium rectale*, and *Faecalibacterium prausnitzii* are known for their critical roles in producing short-chain fatty acids (SCFAs), such as butyrate, which have protective effects against conditions like inflammatory bowel diseases, colorectal cancer, and obesity (Tudela et al., 2021). The high abundance of these keystone species suggests a healthy gut microbiome environment, essential for preventing dysbiosis and maintaining overall gut health (Paine, 1966; Fisher & Mehta, 2014).

### 5. Firmicutes/Bacteroidetes (F/B) Ratio

The Firmicutes/Bacteroidetes (F/B) ratio is a crucial metric for assessing the balance of the two dominant bacterial phyla in the gut microbiome. In this study, the F/B ratio was calculated to be 3.1219, significantly higher than the healthy reference cohort range of 0.14% to 0.76% (Turnbaugh et al., 2009; Cummings et al., 1987; Friedman & Alm, 2012). Firmicutes, which include bacteria beneficial for energy absorption and SCFA production, were more abundant compared to Bacteroidetes, which are involved in breaking down complex molecules like polysaccharides. This higher F/B ratio could indicate a shift in the gut microbiome balance, which may have implications for conditions such as obesity, metabolic syndrome, and other related health issues.

### 6. Condition-Specific Microbial Markers in Gut Health

The analysis identified several microbial markers associated with specific health conditions, such as Irritable Bowel Syndrome (IBS), Inflammatory Bowel Disease (IBD), obesity, cardiovascular diseases, and mental health conditions (Garrett et al., 2010; Scher et al., 2013; Crovesy et al., 2020). These biomarkers are essential for diagnosing and monitoring these conditions, providing valuable insights into the state of an individual’s gut health. For example, alterations in the abundance of specific bacteria were linked to IBS and IBD, while changes in microbial markers were associated with obesity and cardiovascular diseases (Case et al., 2007; Le et al., 2013; Yassour et al., 2016). Identifying these biomarkers is critical for developing personalized treatment plans and advancing personalized medicine in gut health.

## Conclusion

In this study, we introduced PANOMIQ as a comprehensive, end-to-end software solution for the unified analysis of whole-genome, exome, and microbiome data. Unlike traditional tools that are often limited to either primary, secondary, or tertiary analysis, PANOMIQ seamlessly integrates all stages of the analytical process into a single, robust platform. This all-in-one capability not only simplifies the workflow but also eliminates the need for multiple, disparate tools, which can introduce inefficiencies and potential errors.

What truly sets PANOMIQ apart is its ability to deliver rapid, yet comprehensive analysis without compromising on accuracy or depth. While existing solutions that attempt to offer end-to-end processing often sacrifice either speed or robustness, PANOMIQ manages to balance both, enabling researchers and clinicians to process and analyze vast amounts of genomic and microbiome data within hours. This rapid turnaround time is crucial in both research and clinical settings, where timely insights can drive decision-making and advance our understanding of complex biological systems.

By combining speed, comprehensive analysis, and ease of use, PANOMIQ represents a significant advancement in the field of bioinformatics. It empowers researchers to explore the intricate relationships between genomic and microbiome data more efficiently and effectively, paving the way for new discoveries in personalized medicine and disease research. As an end-to-end solution, PANOMIQ not only addresses current challenges but also sets a new standard for future genomic and microbiomic data analysis tools.

## Discussion

The integration of whole-genome, exome, and microbiome data is crucial for advancing our understanding of complex biological systems, yet it remains a significant challenge due to the scale, heterogeneity, and complexity of these datasets. Traditionally, researchers have relied on a combination of tools, each specialized in either primary, secondary, or tertiary analysis, to process and interpret genomic data (Glotov et al., 2023, Wen et al., 2023, Simon et al., 2023). This fragmented approach, while functional, often results in inefficiencies, inconsistencies, and increased potential for errors due to the need to transfer data between different platforms (Pérez-Cobas et al., 2020).

PANOMIQ addresses these limitations by offering a truly unified, end-to-end solution. Unlike other software that typically focuses on a single aspect of the analysis pipeline, PANOMIQ integrates all stages—from raw data processing (primary analysis) and variant calling (secondary analysis) to functional annotation and interpretation (tertiary analysis)—into a single platform. This holistic approach not only streamlines the workflow but also ensures consistency across the entire analytical process, reducing the risk of data loss or misinterpretation that can occur when using multiple tools.

One of the most remarkable features of PANOMIQ is its ability to perform these comprehensive analyses rapidly, often within hours. In many cases, existing end-to-end solutions that attempt to provide similar capabilities are either too slow or compromise on the robustness and depth of analysis to achieve faster processing times. PANOMIQ, however, leverages cutting-edge algorithms and efficient data processing techniques to deliver both speed and reliability, ensuring that users do not have to sacrifice one for the other. This capability is particularly important in clinical settings, where timely insights can be critical for patient care.

Moreover, PANOMIQ’s unified approach to data analysis allows for a more integrated understanding of the interactions between the genome and the microbiome. By analyzing these data types together, PANOMIQ can uncover relationships that might be missed when they are studied in isolation, providing deeper insights into the role of microbiome-host interactions in health and disease. This comprehensive analysis is a significant advantage over other tools that are limited to either genomic or microbiomic data alone.

The robustness of PANOMIQ’s analytical capabilities, combined with its efficiency, makes it a valuable tool for both researchers and clinicians. For researchers, it offers a powerful means of exploring complex biological questions with greater accuracy and speed. For clinicians, PANOMIQ provides a practical solution for incorporating genomic and microbiome data into patient care, enabling more personalized and informed treatment decisions.

While concluding PANOMIQ sets a new standard in genomic and microbiome data analysis by providing a comprehensive, end-to-end solution that is both robust and efficient. It overcomes the limitations of existing tools, offering an integrated approach that enhances the accuracy, speed, and depth of analysis. As the demand for more sophisticated and timely genomic insights continues to grow, PANOMIQ is well-positioned to meet these needs, advancing both research and clinical applications in the field of genomics and bioinformatics.

